# Temporal Window based Feature Extraction Technique for Motor-Imagery EEG Signal Classification

**DOI:** 10.1101/2021.03.19.436144

**Authors:** Rajdeep Chatterjee, Ankita Datta, Debarshi Kumar Sanyal, Swati Banerjee

## Abstract

Electroencephalogram (EEG) based motor-imagery classification is one of the most popular Brain Computer Interface (BCI) research areas due to its portability and low cost. In this paper, we have compared Wavelet Energy-entropy based different prediction models and empirically proven that temporal window based approach in motor-imagery classification provides more consistent and better results than popular filter-bank approach. In order to examine the robustness and stability of the proposed method, we have also employed multiple types of classifiers at the end and found that mix-bagging (bagging ensemble learning with multiple types of learners) technique outsmarts other frequently used classifiers. In our study, BCI Competition II Data-set III has been used with four experimental setup: (a) The whole signal (for each trial) as one segment, (b) The whole signal (for each trial) is divided into non-overlapping segments, (c) The whole signal (for each trial) is divided into overlapping segments, and (d) The filter-bank approach where the whole signal (each trial) is segmented based on different frequency bands. The result obtained from the experiment (c) i.e. 91.43% classification accuracy which outperforms the other approaches not only in this paper but to best of our knowledge it is the highest performance for this dataset so far.

## 1. Introduction

Brain Computer Interface (BCI) technology is a multi-disciplinary domain that involves subjects such as Neuroscience, Digital Signal Processing, Machine Learning etc [1]. Brain rhythms are generated in our brain while performing a mental task. It is also proved that the brain generates rhythms even if a person is in a completely relaxed situation. In a BCI system, the main objective is to acquire specific brain activities and use the patterns (signal) to command a computer in order to perform a particular task. In order to classify these brain patterns, feature extraction and classification both play an indispensable role [2, 3]. Electro-electroencephalogram (EEG) is one of the most widely used recording technique for the motor-imagery classification problem. EEG has its own benefits such as portability with high temporal resolution and affordability over other techniques [4, 5]. Activities such as hand movements, foot movements or any limb movements are performed under the control of our motor cortex region of the brain. While performing these activities, brain rhythms are produced. Even if we think or imagine about limb movements without doing those physically, then also specific activities can be observed in the same brain region and thus they generate similar brain rhythms. This type of thinking or imagination based activities in BCI are known as motor-imagery (MI) [6, 7]. There are different brain rhythms based on brain functional activities: Delta (< 4Hz), Theta (4 − 7Hz), Alpha (7 − 13Hz), Beta (13 − 25Hz), and Gamma (> 25Hz). Sometimes researchers also use Mu (8 − 13Hz) brain signal, which is specifically obtained from the sensorimotor cortex region of our brain. However, the frequency bands are mostly overlapped between Alpha and Mu. As per our understanding the alpha waves generated from sensorimotor cortex region are termed as Mu. After rigorously studying the concept in literature, only Alpha and Beta bands are used for motor-imagery classification problem in this paper [1, 8, 9].

As motor-imagery EEG signals are non-stationary bio-signals, it has its own characteristics. Many researchers have implemented various different feature extraction techniques for motor-imagery classification problem. In the past [10, 11], we have used wavelet based different feature extraction techniques and compared their performances using support vector machine (SVM) and multilayer perceptron (MLP) classifiers. The obtained classification accuracies are 85% and 85.71% respectively. Lemm [12] used the morlet wavelet as features and bayes quadratic as the classifier to achieve 89.29% accuracy. Similarly, Bashar [13] has used multivariate empirical mode decomposition (MEMD) with short time fourier transform (STFT) for feature extraction. He has also used different variants of classifiers but the best performing one is k-nearest neighbor (KNN with cosine distance metric) and obtained 90.71% accuracy.

Previously, we have experimented various suitable combinations of feature extractions and classifiers for EEG signal classification. However, it is observed that a generic framework for EEG signal classification, specifically motor-imagery signals has not yet been reached so far. This very idea has motivated researchers to explore new methods or frameworks in order to improve the overall performance of the classification. In any classification problem, it is always aimed to obtain high accuracy and it involves different steps and techniques to achieve the expected performance. One very important and popular step in that direction is the use of appropriate feature extraction technique for any signal classification. Here, our primary focus is to obtain high level of performance (in terms of accuracy) by identifying the best suited feature-set for a motor-imagery EEG signal classification.

As our last studies shown that wavelet based feature extraction technique is well suited for motor-imagery classification over other commonly used feature extraction techniques. The best practice is to use energy-entropy as features instead of using the actual wavelet coefficients for classification. It is just to reduce the feature-set length. Here, we have decided to use the same energy-entropy wavelet based feature extraction technique but in multiple temporal windows (i.e. non-overlapping overlapping approaches) for a given motor-imagery EEG signal. At this point, we anticipate that our approach will provide us more insights in terms of discriminative information due to the use of multiple temporal segments over the traditional approach for the same trial. Furthermore, the existing best performing techniques are compared with our proposed method in Table 6 at the last.

The paper has been segmented in five different sections. Section 1 is used to introduce the BCI concept, EEG theoretical aspects and related studies. The experimental background is presented in section 2. In section 3 and 4, the the proposed methods and the result & analysis are discussed respectively. Finally, we conclude the paper in section 5.

## 2. Experimental Background

### 2.1. Used Dataset

The used EEG signals i.e. dataset III of BCI competition II (2003) has been provided by the Department of Medical Informatics, Institute for Biomedical Engineering, University of Technology, Graz [14]. The sampling rate of the dataset is recommended at 128*Hz*. The EEG signal was recorded using the standard IEEE 10 20 electrode placement system (Fig. 1). Although EEG signals are available for three different electrodes *C*3, *Cz* and *C*4, we have considered observations from *C*3 & *C*4 electrodes as they are dominant for human left-right hand movements. According to the website [14], dataset contains left-right hand movements of EEG signal from a healthy female subject for 6 seconds as shown in Fig. 2. We selected this dataset instead of other BCI datasets as it has longer available motor-imagery duration for each trials which suits our proposed temporal window approach.

**Figure 1:**
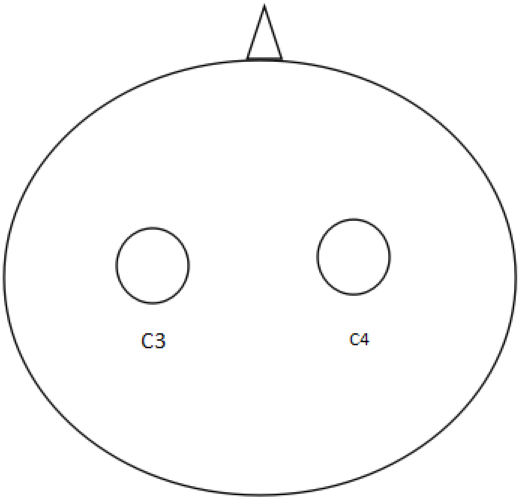
IEEE 10 − 20 electrodes placement for *C*3 and *C*4 electrodes

**Figure 2:**
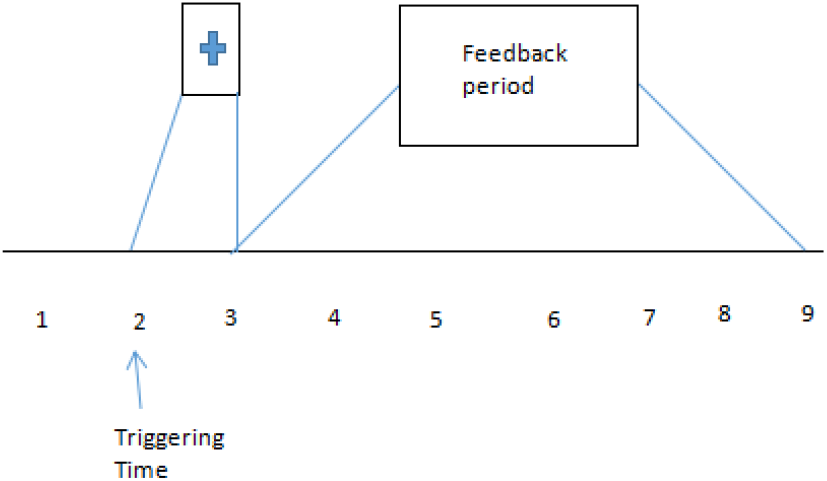
Region of interest (6 sec.) extracted from raw EEG signal for an electrode

**Figure 3:**
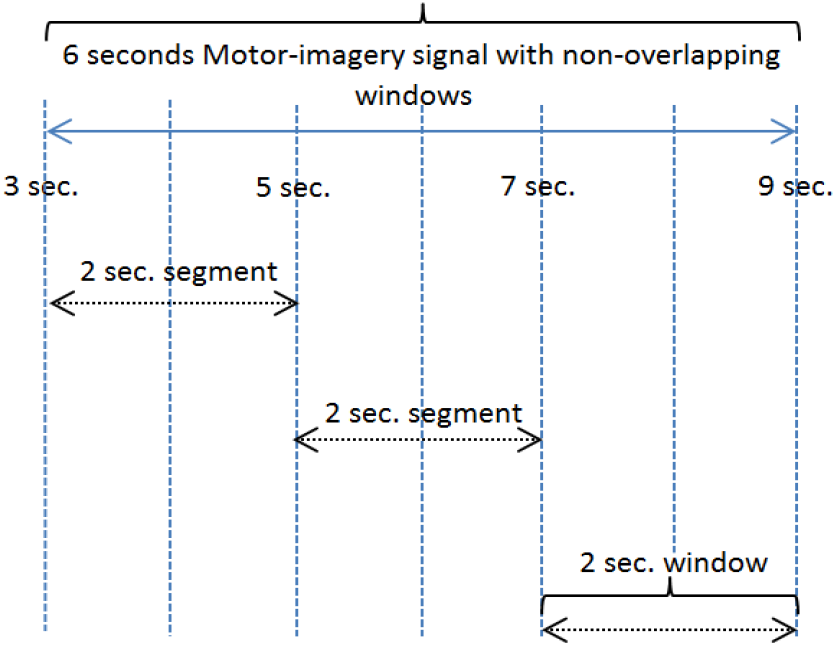
Diagrammatic representation of non-overlapping temporal window approach in Exp-(b)

**Figure 4:**
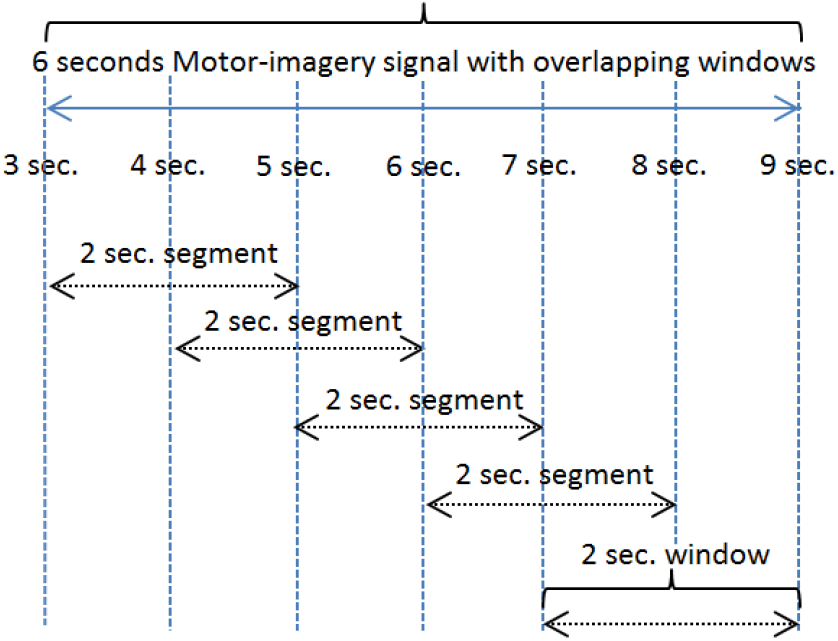
Diagrammatic representation of overlapping temporal window approach in Exp-(c)

**Figure 5:**
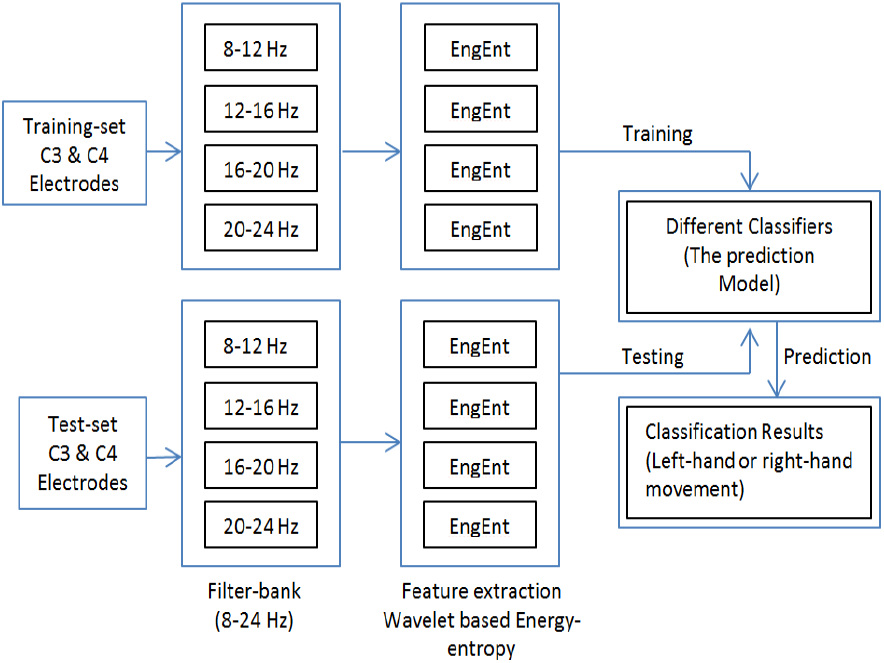
The block diagram of filter-bank approach in Exp-(d)

**Figure 6:**
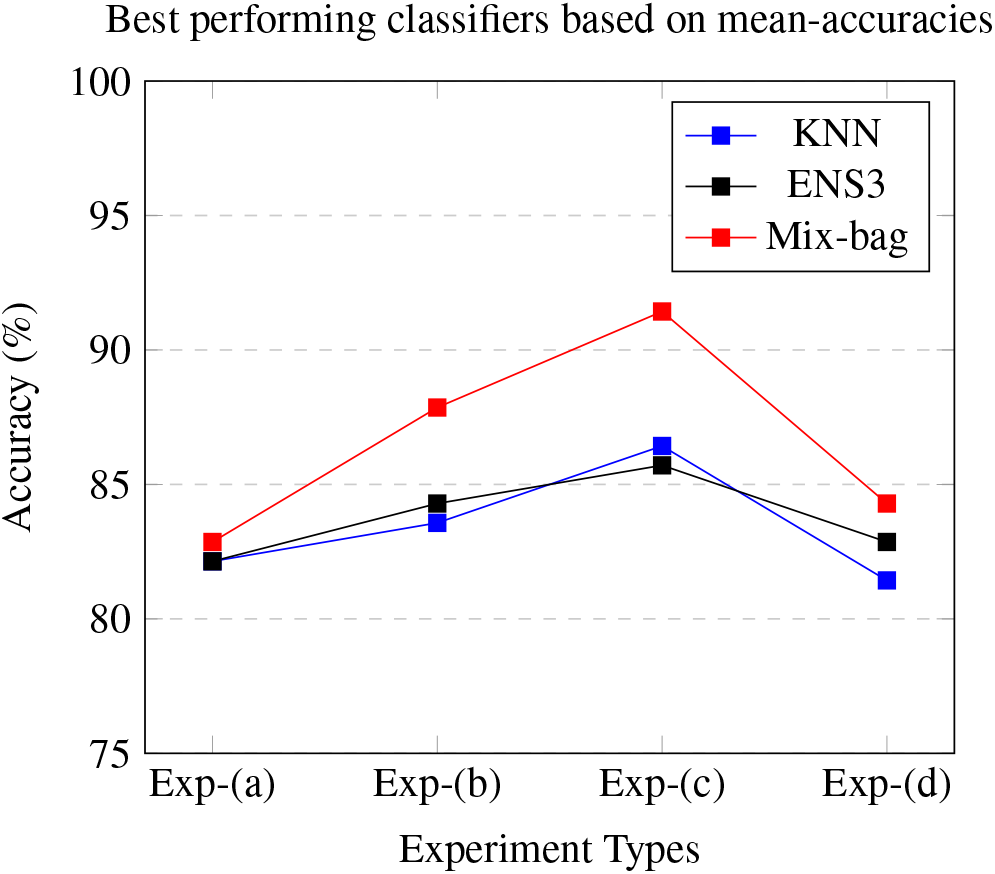
Graphical representation of accuracies obtained from different experiments

**Figure 7:**
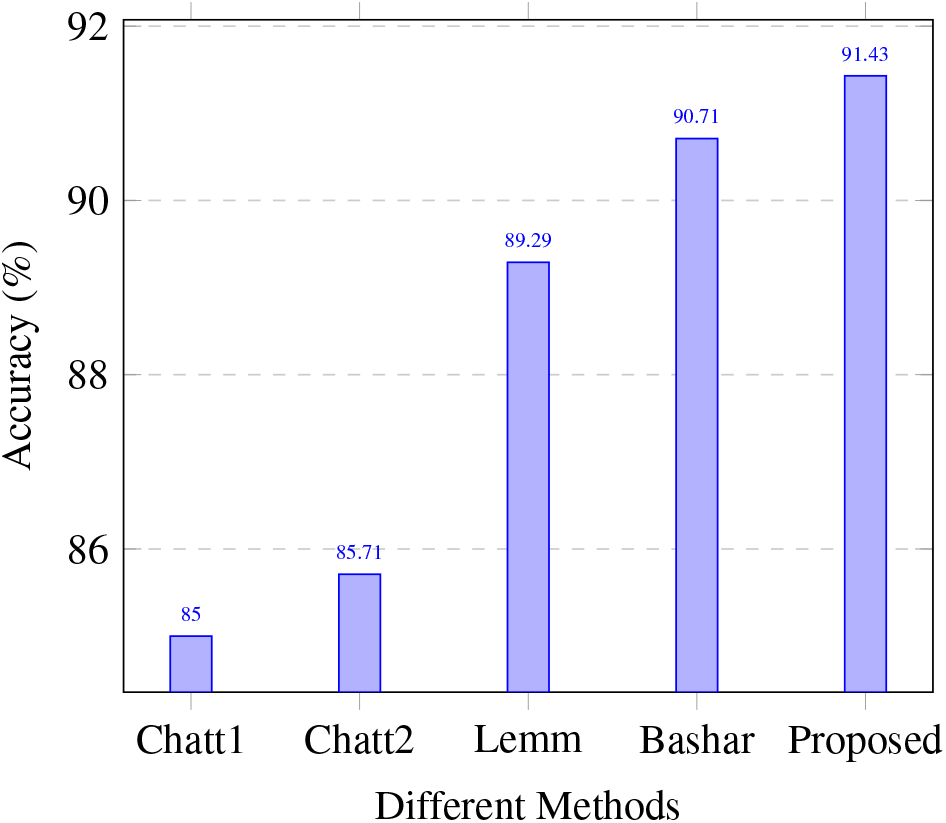
Representation of few best accuracies obtained for BCI Competition II (2003) Data-set III

The desired frequency bands have been filtered from the raw input EEG signal. The whole dataset is of 280 trials (i.e. samples) and a balanced dataset, is divided into 140 trials for training-set and 140 trials for test-set. In order to eliminate the noise from the interested frequency bands, an elliptic bandpass filter, with cut off frequencies of 0.5Hz and 30Hz, has been employed. [7, 10, 11].

### 2.2. Feature Extraction

Wavelet transform seeks to achieve the best trade-off between the temporal and frequency resolution. Instead of using sines and cosines, the wavelet transform uses finite basis functions called wavelets. This single finite-length waveform is known as the mother wavelet. The wavelet transform basically represents the original input signal in the form of linear combination of basis functions. Different varieties of feature extraction techniques are available based on wavelet transform. From our past research work, we have concluded that wavelet based energy-entropy method is well suited for EEG based motor-imagery classifications. [10, 15]

#### Wavelet energy and entropy (EngEnt)

The wavelet based approach is a very suitable method for the feature extraction of EEG signals due to its non-stationary characteristics. It is also proven empirically in literatures as well as in our past works. It can distinguish between both temporal and spectral domain features of a signal. It decomposes the signal into detail and coarse approximation coefficients. The halved output of the first high-pass and low-pass filters provide the detail *D*1 and approximation *A*1 coefficients respectively. This derivation is repeated for another two times to its third level. The Daubechies (db) basis function of order 4 & *D*3 level features have been used in this study [16]. Subsequently, energy and entropy have been computed (Eq. 1 & 2) from the *D*3 featureset instead of using the actual wavelet coefficients as features in order to reduce the dimensionality [10, 16, 17, 18].

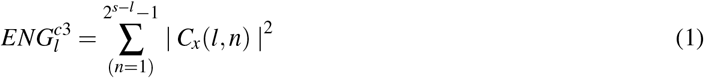

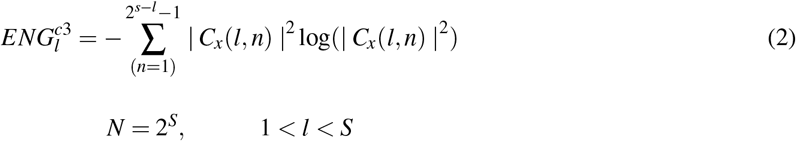

The 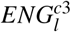 and 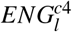 are the computed energy and entropy features for the *l*^*th*^ trial of *C*3 electrode. Similarly, energy & entropy for *C*4 electrode are also obtained. *C_x_*(*l, n*) indicates the input signal of *l*^*th*^ trial and *n* suggests the samples.

### 2.3. Classifiers

In this paper, different types of classifiers have been used: Naive Bayes, K Nearest Neighbor, Support Vector Machine with different kernels and popular boosting & bagging classifiers [19, 20, 21, 22].We have taken the mean of KNN outputs by varying the *K* values from 3 to 35 and the *cosine* distance metric is used. Also, The number of learners in boosting (i.e. ENS1 & ENS2) and the number of bags for bagging ensemble approaches (i.e. ENS3) are averaged by varying them from 10 to 100. It must be mentioned that for all normal ensemble approaches the learner type is always fixed as *decision tree*. Besides these set of classifiers, we have also employed another mixture-bagging (*mix-bag*) ensemble technique in our study borrowed from our past work [23, 24]. The mix-bag ensemble classifier is formed by using multiple types of weak learners (here mix-bag-size=5) for different bags. The mix-bag technique which uses multiple learners demonstrates the best performance due to its higher diversity over the single classifier type. Specifically for bagging and mix-bag, the sample bag sizes are kept at 70% of the actual training dataset. This configuration of parameters are chosen due to its stable and consistent performance after rigorous trials. The details of all the used classifiers are given in Tables 1 & 2.

**Table 1:**
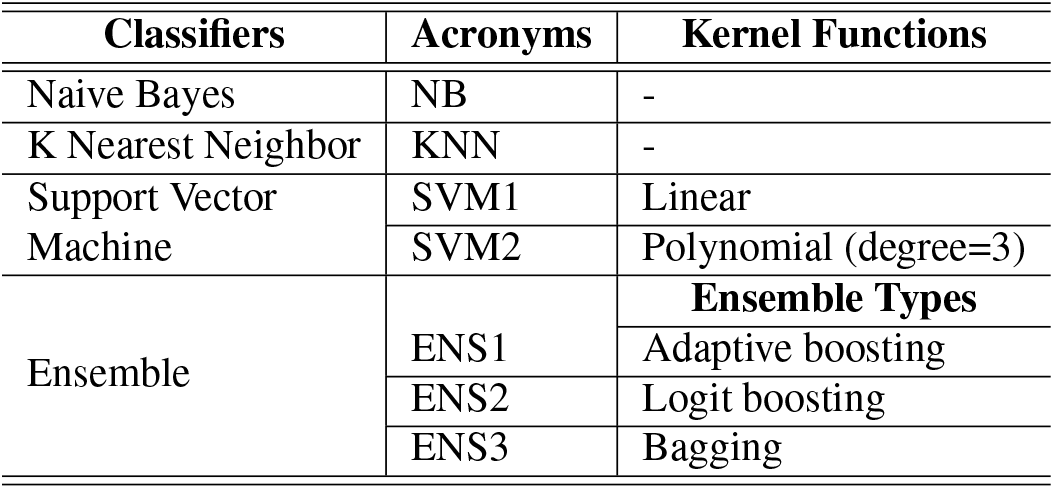
Classifiers configuration used in this paper

**Table 2:**
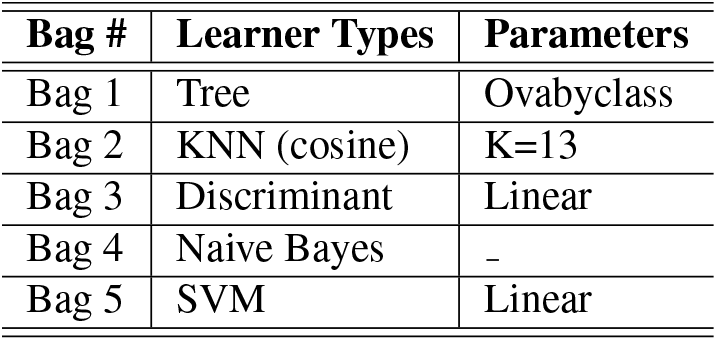
Configuration of the mix-bag ensemble classifier with 5 different learners

| Bag # | Learner Types | Parameters |
| --- | --- | --- |
| Bag 1 | Tree | Ovabyclass |
| Bag 2 | KNN (cosine) | K=13 |
| Bag 3 | Discriminant | Linear |
| Bag 4 | Naive Bayes | - |
| Bag 5 | SVM | Linear |

### 2.4. System Configuration

The experiments are implemented using MATLAB 2016a on an Intel(R) Core(TM) *i*5 − 6200*U* CPU 2.40*GHz* with 8*GB* RAM with 64 bits Windows 10 operating system. The feature extractions and classifications of EEG signals are done by two separate MATLAB scripts.

## 3. The Proposed Approaches

We have discussed the EEG signal types, the preprocessing and wavelet based feature-extraction techniques in the above sections. Also, the theoretical background of basic ensemble learning approaches are also explained precisely. The main objective of this paper is to examine the effects of different feature extraction (overlapping/non-overlapping window based) approaches on EEG based motor-imagery classification. Our initial anticipation before the experiment is that the use of overlapping or non-overlapping feature extraction approach performs better over the use of the traditional feature extraction approach. In order to prove our claim, we have segregated our experiments into four categories and these are presented briefly in Table 3.

**Table 3:** The Brief Description of each Experiment

| Experiments | Description |
| --- | --- |
| **Exp-(a)** | The whole EEG signal (6 Sec.) for each trial is considered as input and wavelet energy-entropy features are extracted from its Alpha & Beta sub-bands. |
| **Exp-(b)** | The whole EEG signal (6 Sec.) for each trial is divided into three *non-overlapping* (2 Sec. each) segments as input and wavelet energy-entropy features are extracted from its Alpha & Beta sub-bands from each segment. (Fig. 3) |
| **Exp-(c)** | The whole EEG signal (6 Sec.) for each trial is divided into five *overlapping* (2 Sec. each) segments as input and wavelet energy-entropy features are extracted from its Alpha & Beta sub-bands from each segment. (Fig. 4) |
| **Exp-(d)** | A Filter-bank with four bands from 8Hz to 24Hz with 4Hz intervals are employed to the whole EEG signal (6 Sec.) for each trial as input and wavelet energy-entropy features are extracted from each band. (Fig. 5) |
| Note: Elliptic bandpass filter is used to extract different frequency bands from the EEG input signal. |  |

As it is shown in Fig. 2 that here each raw EEG signal is of 9 seconds. However, the signal extracted form last 6 seconds is considered for motor-imagery classification. As of now, various authors implemented different feature extraction techniques on this 6 seconds trial considering it as a single entity. It also be noted that we have two 6 seconds signals from two electrodes *C*3 and *C*4. So, the complete 12 seconds signal needs to be extracted in meaningful features for a trial (i.e. single instance). Now, we use energy-entropy wavelet based feature extraction not on the complete 6 seconds signal rather uses multiple windows of 2 seconds to implement it. Our window size is of 2 seconds in order to maintain the uniformity of our experiment (as the original signal is 6 seconds long). The imagination of hand movements (right hand or left hand) based on a 3 seconds cue is actually recorded in this 6 seconds signal per electrode per trial. Such imagination may vary even in that 6 seconds signal and thus a traditional feature extraction approach may not capture the detailed discriminative information if it is applied on the whole signal. on the other hand, if we implement it in multiple overlapping or non-overlapping segments of the original 6 seconds EEG signal, it provides us some additional information which are not captured in earlier cases. Here, our notion is obvious in nature but it also raises question of the suitable number of windows for a signal and its length. In this paper, it is kept at 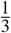 of the original signal. The number of features used for experiments (a), (b), (c) & (d) are 8, 24, 40 and 16 respectively.

## 4. Results and Analysis

The obtained results are shown in Table 4. Three different fundamental types of classifiers have been used with total eight variants. These classifiers are employed over four separate datasets obtained from different experiments (described in Table 3). We have followed the hold-out technique for model building and also executed the process several times independently for KNN and ensemble approaches to have a stable performance. The mean-accuracies for each classifier is also calculated in Table 4 in order to get an overall impression of the best performing classifier. The top three best performing classifiers are Mix-bag, ENS3 and KNN.

**Table 4:** Accuracies (%) obtained from experiments based on different classifiers using EngEnt

| Experiments | NB | KNN | SVM1 | SVM2 | ENS1 | ENS2 | ENS3 | Mix-bag |
| --- | --- | --- | --- | --- | --- | --- | --- | --- |
| Exp-(a) | 78.57 | 82.14 | 78.57 | 76.43 | 81.43 | 82.14 | 82.14 | 82.86 |
| Exp-(b) | 81.43 | 83.57 | 84.29 | 79.29 | 84.29 | 83.57 | 84.29 | 87.86 |
| Exp-(c) | 83.57 | 86.43 | 85.00 | 80.00 | 86.43 | 86.43 | 85.71 | 91.43 |
| Exp-(d) | 75.71 | 81.43 | 81.43 | 78.57 | 77.14 | 76.43 | 82.86 | 84.29 |
| Mean | 79.82 | 83.39 | 82.32 | 78.57 | 82.32 | 82.14 | 83.75 | 86.61 |

As we have already expected earlier that overlapping and non-overlapping based feature extraction with wavelet based energy-entropy provides better and consistent classification accuracies as compared to the popular filter-bank approach and the traditional feature extraction technique i.e. *Exp –* (*a*). Additionally, the overlapping approach gives the best performance over the three other approaches mentioned above. The comparative performance of all the four approaches can be represented as follows:

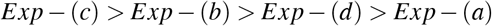

On the other hand, the Mix-bag classifier stands top over all other classifiers due to its inherent diversity introduced by multiple types of learners. The proposed Exp-(c) using Mix-bag classifier gives us highest ever classification accuracy of **91.43%** and a comparative presentation is also given in Table 6 with few other best classification performances for the BCI competition II (2003) dataset III.

The obtained results are validated statistically by using improved Friedman Test [25]. It provides ranks to different algorithms (classifiers) for each separate datasets. The best performing classifier gets the rank 1, subsequent performers get subsequent ranks. In case of ties, average ranks are assigned to those entries. The Friedman test compares the average ranks of algorithms i.e. *R_j_*.

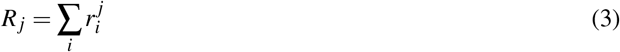

where 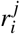 is the rank of the *j^th^* of *k* classifiers on the *i^th^* of *N* datasets. It works under the null hypothesis that all classifiers are having equal rank. It means all classifiers are equivalent in performance. The Friedman statistic:

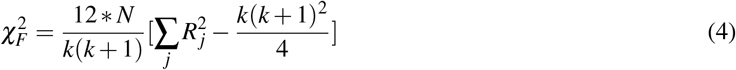

where *N* is the number of datasets, *k* is the number of algorithms (classifiers) and 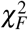 distribution with *k* − 1 degree of freedom is followed in Friedman test. However, it was proven that Friedman test is a conservative statistic and in the year 1980, Davenport derived an improved statistic *F_F_* [25]. The *F_F_* follows F-distribution with *k* − 1 and (*k* − 1)(*N* − 1) degrees of freedom.

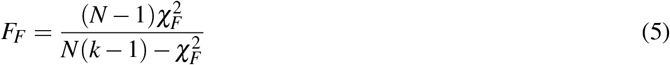

Now using the values of Table 5, the Eq. 4 and Eq. 5 can be re-written as:

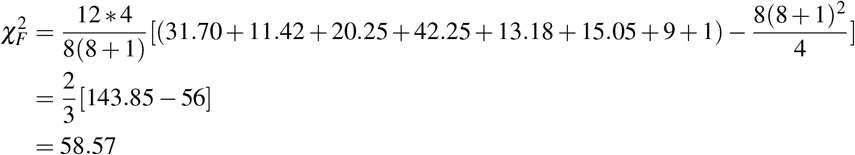

**Table 5:** Rank based comparison of different classifiers and obtain average rank *R_j_* & 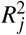

| Experiments | NB | KNN | SVM1 | SVM2 | ENS1 | ENS2 | ENS3 | Mix-bag |
| --- | --- | --- | --- | --- | --- | --- | --- | --- |
| Exp-(a) | 5.5 | 3 | 5.5 | 6 | 4 | 3 | 3 | 1 |
| Exp-(b) | 5 | 4.5 | 3 | 6 | 3 | 4.5 | 3 | 1 |
| Exp-(c) | 6 | 3 | 5 | 7 | 3 | 3 | 4 | 1 |
| Exp-(d) | 6 | 3 | 4.5 | 7 | 4.5 | 5 | 2 | 1 |
| avg. rank, $R_j$ | 5.63 | 3.38 | 4.5 | 6.5 | 3.63 | 3.88 | 3 | 1 |
| $R_j^2$ (Eq. 4) | 31.70 | 11.42 | 20.25 | 42.25 | 13.18 | 15.05 | 9 | 1 |

**Table 6:**
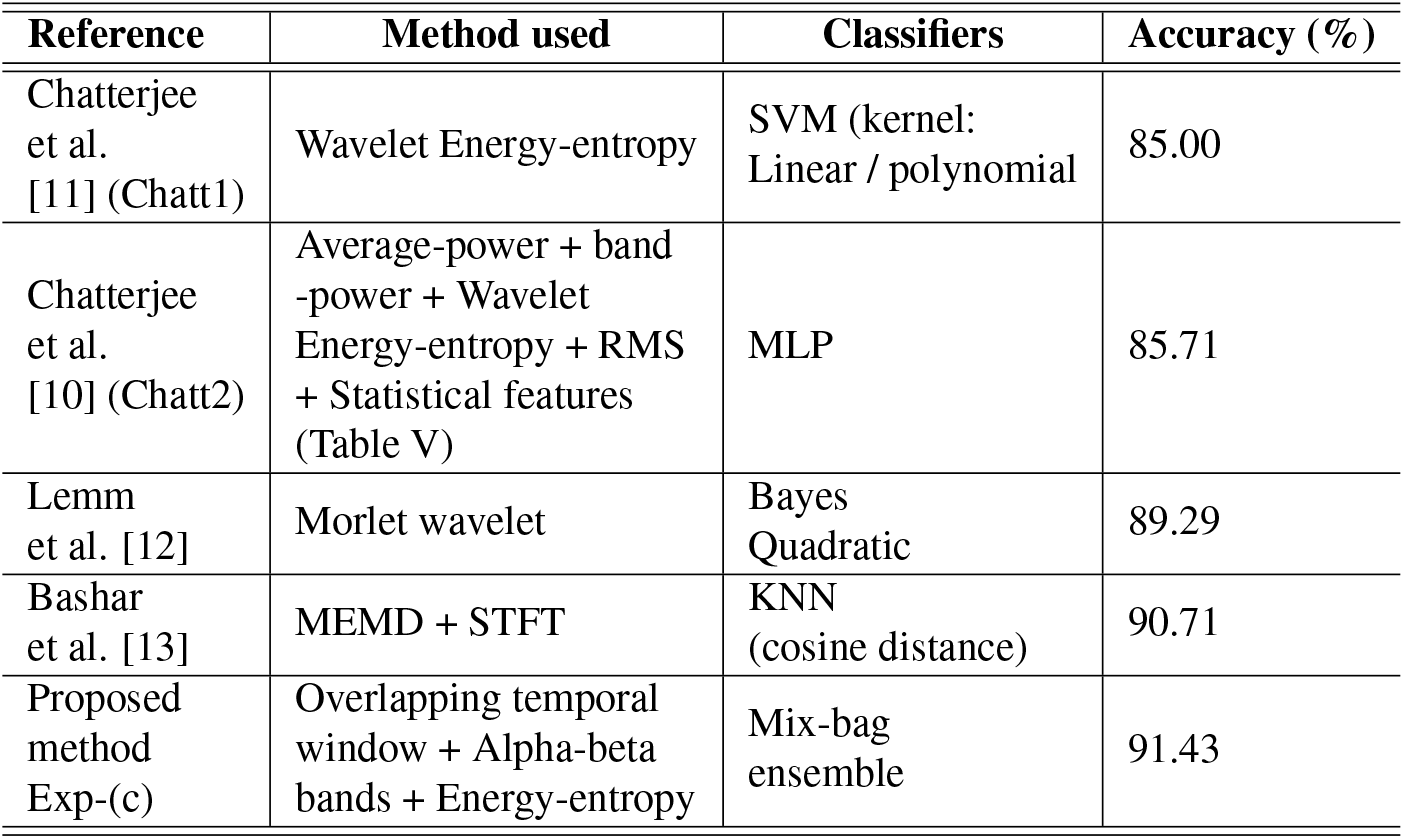
Comparative analysis of few best performing techniques on BCI 2003 Data-set III

| Reference | Method used | Classifiers | Accuracy (%) |
| --- | --- | --- | --- |
| Chatterjee et al. [11] (Chatt1) | Wavelet Energy-entropy | SVM (kernel: Linear / polynomial) | 85.00 |
| Chatterjee et al. [10] (Chatt2) | Average-power + band -power + Wavelet Energy-entropy + RMS + Statistical features (Table V) | MLP | 85.71 |
| Lemm et al. [12] | Morlet wavelet | Bayes Quadratic | 89.29 |
| Bashar et al. [13] | MEMD + STFT | KNN (cosine distance) | 90.71 |
| Proposed method Exp-(c) | Overlapping temporal window + Alpha-beta bands + Energy-entropy | Mix-bag ensemble | 91.43 |

The critical value of 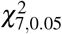 is 14.07. Similarly,

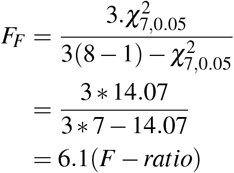

The critical value of *F*_7,21_-distribution for *α*=0.05 is 2.49. In both the statistical tests, the critical values obtained in our research are less than the actual ratios. So, the null hypothesis i.e. all classifiers are equivalent, is incorrect. The *F – ratio* is likely to occur by chance with a *p <* 0.05. Therefore, mix-bag is correctly ranked 1 in our study.

## 5. Conclusion

Physiologically, similar brain functioning occurs for the same motor-imagery movements. However, the discriminative information varies with time for each trial even for the same subject. Here the 6 seconds long motor-imagery EEG signal contains the discrimination event which is captured well in overlapping temporal window based feature extraction approach over other alternatives. From our empirical results, it is suggested that more use of time segments (windows) provide more insight to the discriminants for the same input signal. However there is a trade-off between the number of time windows used and the quality of classification. It increases the number of features as the number of overlapping windows goes up. It is up to the researchers to decide his/her priority whether he/she prefers reasonable good classification accuracy with moderate features or goes for very high performance. Thus, it propels researchers to explore it further to attain higher performance than ever before. In future, our focus will be on the selection of an adaptive window length for the motor-imagery EEG signal classification using more datasets obtained from different healthy subjects.

